# Native ultrastructure of fresh human brain vitrified directly from autopsy revealed by cryo-electron tomography with cryo-plasma focused ion beam milling

**DOI:** 10.1101/2023.09.13.557623

**Authors:** Benjamin C. Creekmore, Kathryn Kixmoeller, Ben E. Black, Edward B. Lee, Yi-Wei Chang

**Affiliations:** Translational Neuropathology Research Laboratory, Department of Pathology and Laboratory Medicine, Perelman School of Medicine, University of Pennsylvania, PA, USA; Department of Biochemistry and Biophysics, Perelman School of Medicine, University of Pennsylvania, PA, USA; Biochemistry and Molecular Biophysics Graduate Group, Perelman School of Medicine, University of Pennsylvania, PA, USA; Institute of Structural Biology, Perelman School of Medicine, University of Pennsylvania, Philadelphia, PA, USA

## Abstract

Ultrastructure of human brain tissue has traditionally been examined using electron microscopy (EM) following chemical fixation, staining, and mechanical sectioning, which limit attainable resolution and introduce artifacts. Alternatively, cryo-electron tomography (cryo-ET) offers the potential to image unfixed cellular samples at higher resolution while preserving their native structures, but it requires samples to be frozen free from crystalline ice and thin enough to image via transmission EM. Due to these requirements, cryo-ET has yet to be employed to investigate the native ultrastructure of unfixed, never previously frozen human brain tissue. Here we present a method for generating lamellae in human brain tissue obtained at time of autopsy that can be imaged via cryo-ET. We vitrify the tissue directly on cryo-EM grids via plunge-freezing, as opposed to high pressure freezing which is generally used for thick samples. Following vitrification, we use xenon plasma focused ion beam (FIB) milling to generate lamellae directly on-grid. In comparison to gallium FIB, which is commonly used for biological samples, xenon plasma FIB is powerful enough to efficiently mill large volume samples, such as human brain tissue. Additionally, our approach allows for lamellae to be generated at variable depth inside the tissue as opposed to being limited to starting at the surface of the tissue. Lamellae generated in Alzheimer’s disease brain tissue and imaged by cryo-ET reveal intact subcellular structures including components of autophagy and potential tau fibrils. Furthermore, we visualize myelin revealing intact compact myelin and functional cytoplasmic expansions such as cytoplasmic channels and the inner tongue. From these images we also measure the dimensions of myelin membranes, providing insight into how myelin basic protein forces out oligodendrocyte cytoplasm to form compact myelin and tightly links intracellular polar head groups of the oligodendrocyte plasma membrane. This approach provides a first view of unfixed, never previously frozen human brain tissue prepared by cryo-plasma FIB milling and imaged at high resolution by cryo-ET.

## Introduction

Transmission electron microscopy (transmission EM) has long been used to inspect cellular structures that are not readily visible or distinguishable via light microscopy. While allowing relatively high-resolution imaging, this technique historically applies chemical fixation, dehydration, and heavy element staining to the sample to ensure adequate visualization of these structures. These preparation methods can introduce damage or artifacts that can perturb the native architecture of cellular components, especially delicate structures such as cytoplasmic channels within myelin [7, 15]. Preservation of native cellular architecture is improved using high pressure freezing (HPF) followed by freeze substitution, but these approaches still involve organic solvent incubation, resin embedding, and mechanical thin-sectioning of the sample for EM visualization, which also reduce the attainable resolution [23, 30, 34].

Excitingly, the establishment of cryogenic transmission electron microscopy (cryo-EM) has eliminated the requirement of sample fixation or staining for structural visualization. This allows for better preservation of native structures, enabling the visualization of details not attainable by traditional EM methods. In the field of neurodegeneration, cryo-EM has mostly been performed on purified protein samples, such as resolving to near-atomic resolution the structures of aggregates extracted from diseased brains including tau, amyloid-β, α-synuclein, and TDP-43 [1, 31, 35, 36]. However, these aggregates have all been removed from their native environment and thus do not provide a complete understanding of the associated cellular processes. For phenomena difficult to isolate or recapitulate *in vitro*, such as vesicular architecture and myelination, traditional EM methods remain the most informative to date.

Bridging the benefits of both traditional EM imaging and *in vitro* cryo-EM, cryo-electron tomography (cryo-ET) has emerged as a technique for using cryo-EM to image cells and tissues to better understand the context and three-dimensional organization of structures of interest. In cryo-ET, tilt series are collected at regions of interest by physically tilting a sample and taking images at different angles. These images are then reconstructed into three-dimensional tomograms. Importantly, to attain high resolution, cryo-ET, and cryo-EM in general, relies on adequate vitrification to avoid perturbation of structures by crystalline ice and also requires samples thin enough to allow electrons through the sample for transmission EM. For purified aggregates or thin cells in culture, this is relatively easily achievable. Some phenomena, like neurodegenerative diseases and myelination, however, are not able to be fully reconstituted in cell models therefore necessitate the use of tissue.

For thin samples, rapid vitrification to preserve native architecture is commonly achieved via plunge-freezing sample into liquid ethane or ethane/propane mixture directly on a cryo-EM grid. However, plunge-freezing alone is only effective to less than about 10 μm thickness [18]. HPF has been used to fully vitrify thicker samples, such as tissues [34]. HPF, however, is low throughput and makes sample thinning prior to cryo-ET challenging. In order to allow for sample thinning directly on a cryo-EM grid, similar to what can be done with samples that are plunge-frozen, components of the high pressure freezer need to be modified or generated [12]. Most approaches using HPF for cryo-ET, therefore, instead employ either cryo-sectioning that leaves significant surface artifacts and distortions [10] or cryo-liftout which is very low throughput and technically challenging [19]. Recently, to avoid these HPF-associated challenges, it was shown that, with the addition of glycerol, plunge-freezing can vitrify some samples up to about 200 μm thick [2], but this may be sample-dependent (described below).

Once vitrified, cryo-focused ion beam (cryo-FIB) milling is currently the method of choice for thinning cryo-EM samples because it introduces many fewer artifacts compared to cryo-sectioning [10, 20]. Gallium is the ion source most commonly used for cryo-FIB milling of biological samples. However, gallium lacks the power to efficiently remove large amounts of material and over time can alter sample architecture due to the depth of gallium interaction with samples [17]. In contrast, xenon plasma-based cryo-FIB milling, primarily used in material science, is capable of efficient thinning of very large samples and has reduced sample interaction depth compared to gallium [4, 13, 39].

Motivated by the aforementioned technical challenges and that conventional tissue preservation methods at time of autopsy (chemical fixation or −80 °C storage) introduce artifacts or damage, we sought to develop a new approach that permits cryo-ET imaging of human brain tissue obtained at time of autopsy without prior fixation or freezing. We describe a protocol for vitreous plunge-freezing of human brain tissue at least 180 μm thick. We further present a method for using xenon plasma-based cryo-FIB milling at currents never previously used in a biological context to generate lamellae suitable for cryo-ET imaging. These lamellae are generated directly on cryo-EM grids at variable depths into the tissue and are not limited to starting at the tissue surface. We demonstrate the capability of this approach by providing examples of the visualization of native subcellular compartments, disease-associated tau aggregates, and the intricate myelin architecture – including insights into the structural organization of myelin basic protein.

## Results

In order to image human brain tissue in its native condition in the absence of freezing or fixation artifacts, we obtained unfixed, never previously frozen samples from the middle frontal cortex of several individuals at time of autopsy (box in Fig. 1a; Fig 1b). Gray matter (box in Fig. 1b) was further cut into small pieces <3 mm in width to approximate the dimensions of a standard cryo-EM grid. Tissue was embedded in cool low-melting point agarose (Fig. 1c) and then vibratome sectioned to a final thickness of about 100 μm (Fig. 1d). Due to the malleable and soft consistency of fresh brain tissue, sectioning thinner than roughly 100 μm was difficult.

**Fig. 1.**
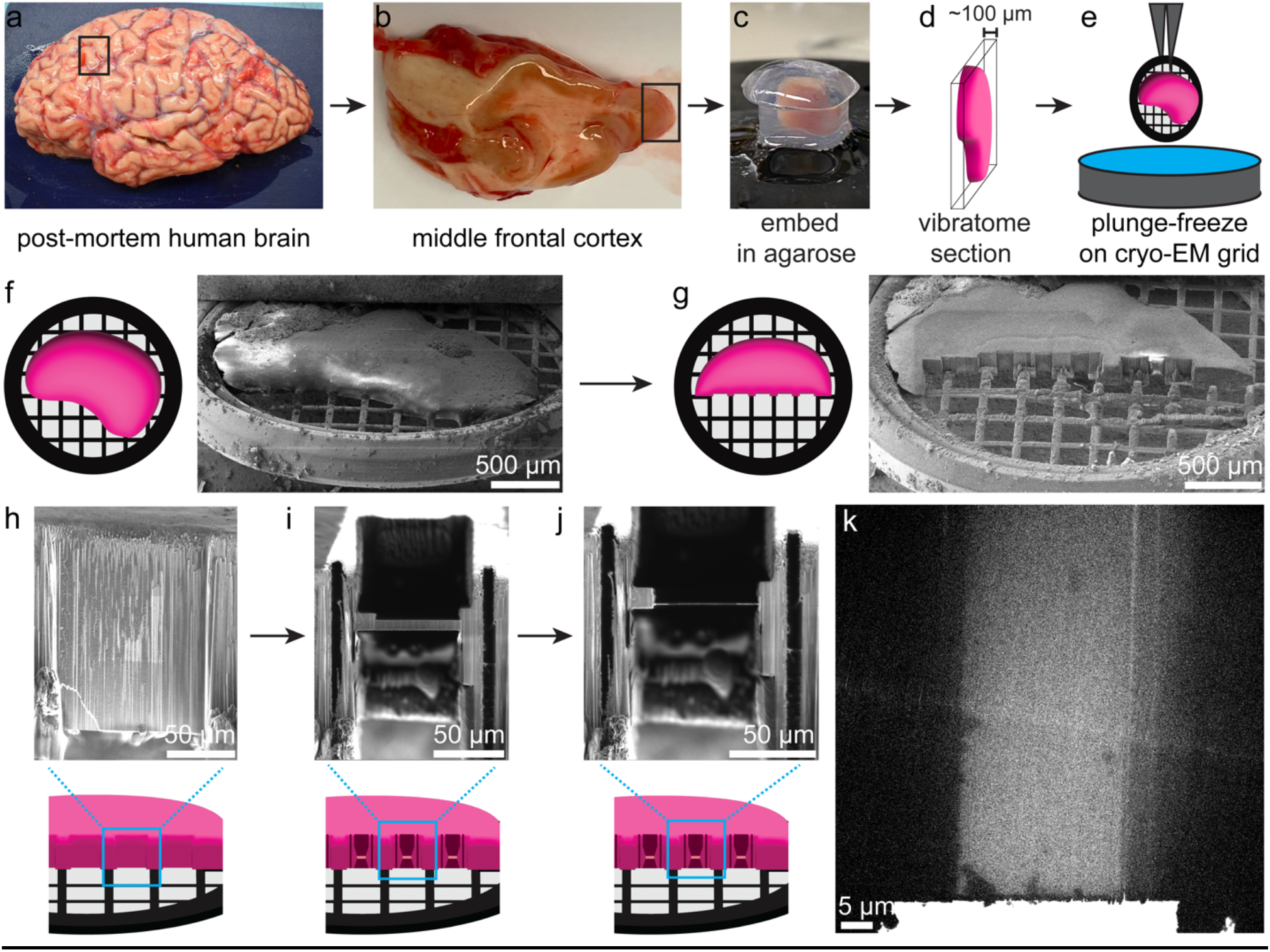
Vitreous cryo-preservation and cryo-FIB milling of fresh, unfixed, never previously frozen post-mortem human brain tissue. **a-e** Schematic of acquisition, sectioning, and freezing of post-mortem human brain tissue showing the procession from gross sample (**a**) used for initial acquisition of middle frontal cortex (black box in **a, b**) to the separation of small segments of cortical gray matter (black box in **b**) from white matter to embedding gray matter sections in 6% low melting point agarose molds (**c**) prior to thin vibratome sectioning (**d**) and incubation in cryoprotectant to plunge-freezing sample into liquid ethane/propane (**e**). **f** Schematic and example SEM image of tissue on a cryo-EM grid within the FIB/SEM prior to milling. **g** Schematic and example SEM image of tissue after trench milling by cryo-FIB. **h-j** Schematic and example FIB images of on-grid lamella milling of tissue to generate final lamellae that are about 400 nm thick (**j**). **k** Example transmission EM image of final lamella suitable for cryo-ET.

We sought to determine whether unfixed, never previously frozen human brain tissue could be vitrified via plunge-freezing on standard EM grids into a liquid ethane/propane mixture (Fig. 1e), since this approach would allow for high-throughput on-grid vitrification and simpler on-grid lamella generation as opposed to the technically-challenging cryo-liftout [28], ultimately making this approach more widely accessible. We first attempted to vitrify tissue by plunge-freezing after an incubation in a cryoprotectant solution of 10% glycerol for 15 min following recent protocols [2]. However, after cryo-FIB milling of these samples (see detailed method below), cryo-EM imaging revealed that the lamellae were not vitreous and images contained Bragg reflections caused by diffraction of the electron beam due crystalline ice (pink arrows Fig. S1a). We next increased the glycerol to 20% and added 1M trehalose based on the ability of trehalose to augment glycerol cryoprotection of stem cells [37]. Tissue was successfully vitrified with this method (Fig. S1b-e). We have successfully used this method with four separate autopsy cases with incubation times down to 70 min for samples 180 μm thick. We do not rule out that shorter incubations in the cryoprotectant solution or thicker tissue sections may also allow full vitrification.

To date, a significant obstacle in FIB milling of large samples has been the substantial amount of time necessary to remove large areas of material. Biological samples have almost exclusively been milled on gallium-based FIBs, but these instruments are limited in their milling rates and consequently are not suitable for handling large tissue samples. We therefore established a Tescan S8000X plasma FIB (PFIB)/Scanning Electron Microscope (SEM) with Leica VCT and VCM for cryo-FIB milling using xenon plasma. FIB current is the rate at which ions are projected at a sample and current is generally proportional to milling rate for a given ion source. Xenon PFIB can achieve higher currents and has a higher milling rate at a given current for most materials when compared to gallium FIB [6]. Xenon PFIB therefore can remove material faster than gallium FIB. Cryo-PFIB has only recently been used for biological samples and in those cases used only beam currents also accessible via gallium FIB [4].

In addition to the obstacles posed by FIB milling rates, large samples, especially those that are thick and cover most of an EM grid surface, pose a geometrical challenge for lamella generation and placement. Considerations for lamella placement include stability of the site as well as avoiding areas of potential mechanical damage from the earlier vibratome sectioning. It must also be feasible to remove all material, including metal grid bars, from above and below the final lamella position since any remaining material will impede the subsequent EM imaging. Because cryo-ET requires sample tilting, even material in front of or behind a lamella can impede imaging when the sample is tilted if not adequately removed. Recent methods have allowed for on-grid lamella generation in relatively large samples [2, 12]. However, like most previous methods, they primarily generate lamellae beginning at the surface of the specimen and so the depth into a specimen is determined by the length and angle of the lamella or a prominent sample edge. For cells and organisms such as *Caenorhabditis elegans* that can be vitrified intact [12], starting a lamella at the surface has minimal disadvantages. However, bulk tissue samples, like human brain, require pre-processing including the use of scalpels, razor blades, and vibratome blades to cut the tissue prior to vitrification. This may result in mechanical damage of the specimen surface. Therefore, we sought to develop a protocol that would allow lamellae to be initiated from the interior of the tissue and thereby avoid the potential tissue disruption and artifacts present at the tissue’s pre-cut surfaces and allow more flexibility in lamella placement.

First, the entire surface of the tissue is coated with a protective layer of organometallic platinum from the PFIB’s gas injection system (Fig. 1f). Next, large amounts of tissue are removed in order to expose an internal tissue face (Fig. 1f-g). This initial ‘trench milling’ is performed with the FIB oriented roughly orthogonal to the plane of the grid and using a beam current of 1 μA, which is not achievable on currently available gallium FIB systems. Roughly 80 μm short of the desired lamella-milling position, the current was lowered to 100 nA for the remainder of the trench milling step. This reduction of the beam current serves to prevent damage of the tissue that will eventually be imaged due to the wide gaussian tail associated with the powerful 1 μA FIB beam. The 100 nA beam current, while available on gallium FIB systems, is better focused when using xenon PFIB [32]. The 100 nA beam is also used to remove grid bars near the internal tissue face, thereby preventing the bars from occluding imaging of final lamellae. Finally, a 10 nA beam is used with a directional polishing pattern to smooth the front of the internal tissue face to prepare this front, newly exposed surface for platinum coating and subsequent lamella generation (Fig. 1g).

Once this trench milling step is completed, the sample is rotated to point the gas injection system needle directly at the newly exposed internal tissue face. The tissue face is iteratively coated with organometallic platinum and then the deposited platinum layer is cured by exposure to the ion beam – an approach that compacts the platinum layer for better protection of the underlying tissue. After platinum coating, lamellae are milled into the tissue face using an approach similar to other on-grid lamella generation methods, starting at 10 nA beam current and gradually reducing to 50 pA for final polishing (Fig. 1h-j) [2, 4, 12]. Expansion joints about 8 μm wide are also milled on both sides of the lamellae. Expansion joints serve to release mechanical stress due to slight movements or fluctuations in temperature thus help preserve lamellae during sample transfer steps (Fig. 1i-j). Lamellae were milled to a final width of about 30-40 μm and a length of about 25 μm to more than 70 μm (Fig. 1j-k). We noticed a reduction in successful lamella completion and survival through sample transfer steps when lamellae were wider than 50 μm.

After milling, grids were transferred to a Titan Krios cryo-electron microscope for imaging. We found that lamellae prepared by the above method could survive the transfer from the PFIB to Titan Krios and are generally also thin enough for cryo-EM imaging (Fig. 1k). Two-dimensional projection images of lamellae at various magnifications (Fig. 1k; Fig. 2a-c; Fig. S2) and tilt angles (Fig. S1b-e) showed no evidence of crystalline ice or gross damage within the lamella and revealed intact subcellular structures. Furthermore, two-dimensional montage imaging at x19,500 magnification provided an overview of the contents of the lamellae and gave important context for individual observations (Fig. 2a). These images reveal the cellular and sub-cellular milieu of unfixed, cryo-preserved human brain tissue. Importantly, many delicate structures often disrupted by conventional autopsy sample freezing/fixation or by mechanical cutting were found to be intact and well-preserved in our lamellae. For example, both myelin and lipid membranes showed no signs of fracture and vesicles maintained their rounded shape (Fig. 2b-c). From two-dimensional images, subcellular structures could be identified in their native environment including endosomes, autophagic vesicles, and lysosomes (Fig. 2a-c; Fig. S2a-b). One striking feature evident throughout our lamellae was the repetitive layering of myelin surrounding axons (Fig. 2a, c; Fig. S2). The repeating unit of myelin, consisting of two lipid bilayer membranes that appears as three distinct lines was preserved in our images, previously described only by fixed and stained tissue [11].

**Fig. 2.**
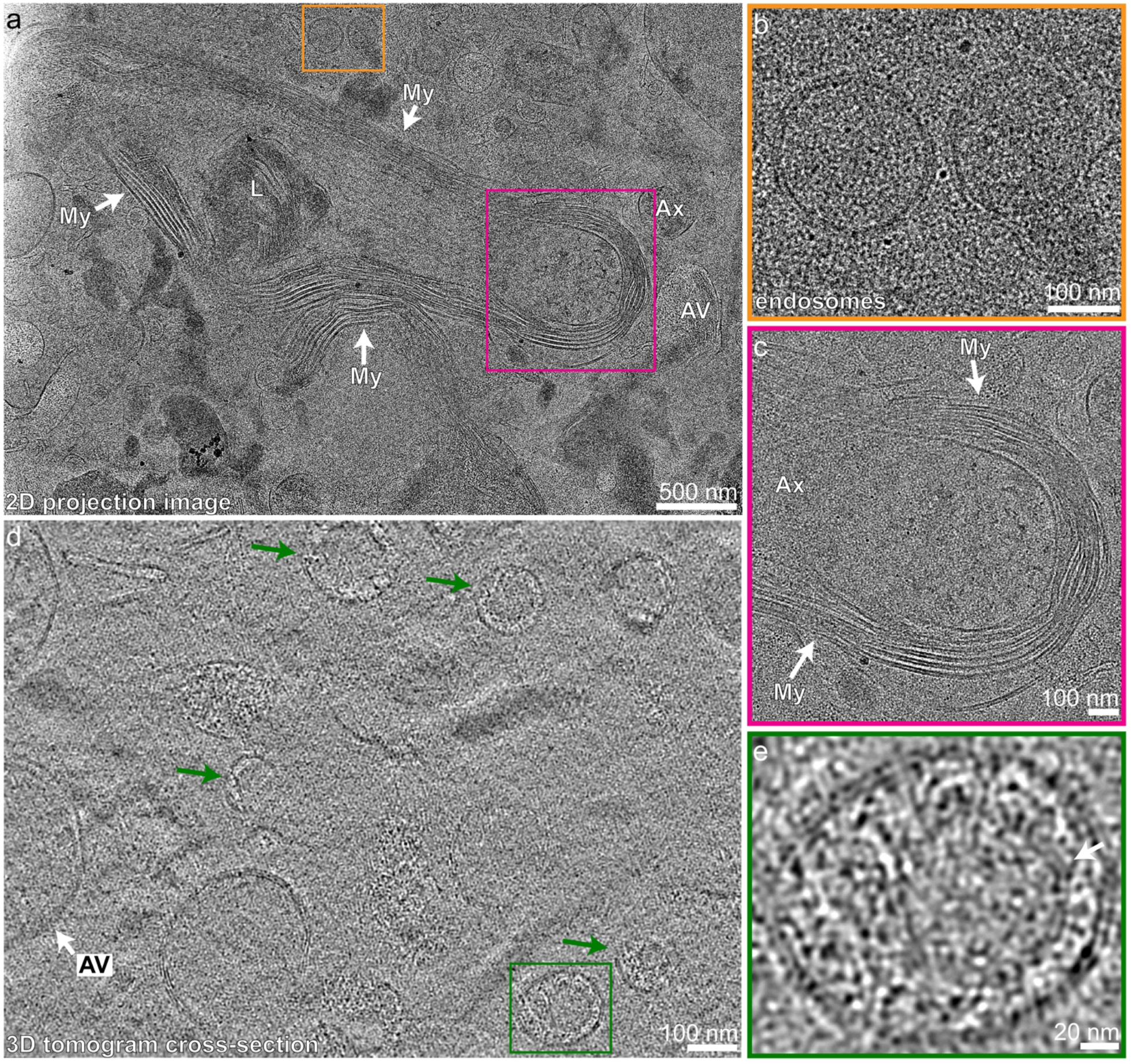
Architecture of cellular and subcellular features is preserved as seen by cryo-EM imaging of lamellae. **a** Two-dimensional projection image at x19,500 of an example lamella that shows preserved architecture of a lysosome (L), unmyelinated axon (Ax), autophagic vesicle (AV), and myelin (My). The orange box is expanded in **b** showing preserved membrane architecture of two putative endosomes. The pink box is expanded in **c** showing a closer view of preserved myelin architecture around a myelinated axon. **d** Cross-section of a denoised tomogram showing an autophagic vesicle (AV) and granular vesicles (green arrows). The green box is expanded in **e** showing a closer view of a granular vesicle with a potential inner membrane (arrow).

In addition to these two-dimensional images, tilt series were acquired on lamellae. The tilt series were then reconstructed into tomograms to reveal the three-dimensional organization of cellular and subcellular structures (Fig. 2d-e; Fig. 3-4; Online Resource 1-4). The three-dimensional data of tomograms also provided better visualization of subcellular membranous structures, such as granular vesicles and phagophore-like structures (Fig. 2d-e; Fig. 3a-e; Online Resource 1-2). We also observed relatively large single-membrane bound vesicles containing heterogeneous cellular elements that may be autophagic vesicles, autophagolysosomes, or lysosomes, an example of which is shown in Fig. 3a (AV with white arrow). Numerous granular vesicles of various sizes were visualized and are composed of an outer membrane surrounding a relatively electron dense spherical core (Fig. 2d-e). In some examples, we observe a tentative, additional layer of membrane surrounding the dense core (Fig. 2e, arrow). The exact identity of these granular vesicles remains to be identified. However, similar granular vesicles of the central nervous system reported at low resolution by traditional EM are thought to contain neuropeptide [25]. In addition, we found crescent shaped cross-sections of single-membrane structures which are narrow in the center and dilated at the ends (Fig. 3a, b-e; Online Resource 2). We observe different forms of these structures that we hypothesize to be different stages of biogenesis (Fig. 3b-e). The peripheral dilation becomes more pronounced as the structure expands and curves inward (Fig. 3b-e). This process seems to mimic phagophore biogenesis observed in yeast and cultured human cells, where there is a dilation of the rim of the forming phagophore [5, 38]. These similarities suggest that the crescent-shaped features are phagophore-like structures. Future work may be able to confirm the identity of these structures as phagophores and shed light on organelle biogenesis directly in human tissue.

**Fig. 3.**
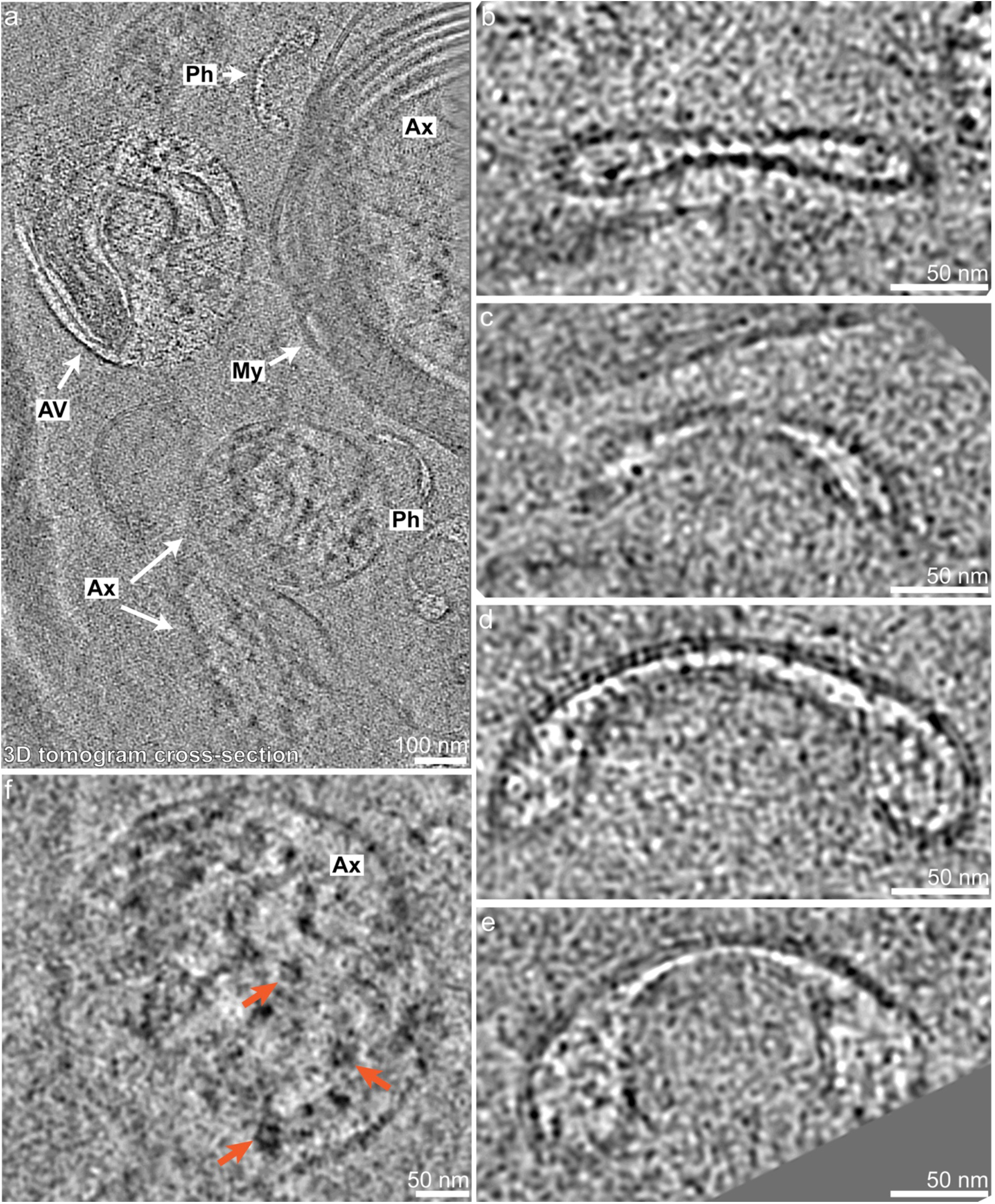
Various subcellular features can be identified in the diverse cellular milieu captured by lamellae in human brain tissue. **a** Cross-section of a denoised tomogram showing an autophagic vesicle (AV) containing various cellular structures, areas of phagophore-like structures (Ph), two unmyelinated axons (Ax), and one larger myelinated (My) axon (Ax). **b-e** Images of phagophore-like structures from the same tomogram shown in **a** that show a single-membrane structure with a narrow center and dilated ends. The axons in **a** contain densities of proteins that run parallel to the axon length. **f** shows a closer look at some of these protein densities including densities consistent with microtubules (orange arrow) that are important for axonal transport.

**Fig. 4.**
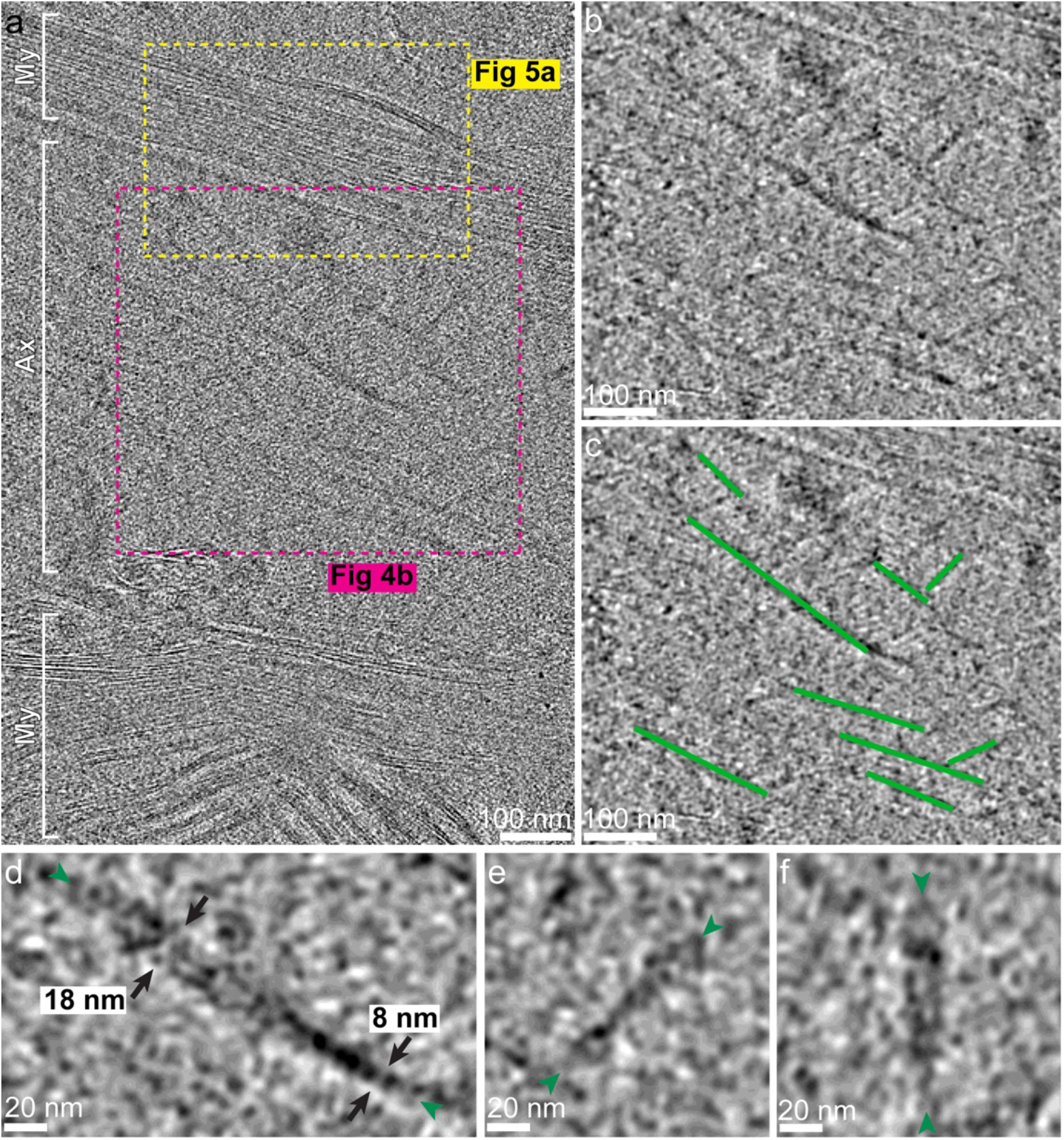
Alzheimer’s disease tau can be visualized and mapped within tomograms. **a** Cross-section of a not denoised tomogram showing a longitudinal view of a myelinated axon containing twisting fibrils consistent with tau fibrils. Myelin (My) surrounds the axon (Ax) on either side in this longitudinal cross-section. **b** The pink box from **a** is shown with denoising to highlight the twisting tau fibrils. **c** Tracings of the fibrils seen in **a** and **b** overlaid with the image from **b. d-f** Closer views of the tau fibrils (ends shown with green arrowheads) with the approximate maximum and minimum width shown with black arrows in **d**.

Beyond simple intracellular membranous structures, both myelinated and unmyelinated axons were visible in several tomograms as transverse, longitudinal, or oblique cross-sections. Axon cross-sections revealed numerous densities that run parallel to the length of the axons which, based on their dimensions and morphology, appear to be microtubules that are abundant in axons due to their roles in maintaining axonal structure and supporting axonal transport (Fig. 3f, orange arrows). Other densities were observed within axons which run parallel to the axon but do not resemble microtubules. For example, Figure 4 shows a longitudinal cross-section of a myelinated axon that contains several long twisting filamentous structures. The filaments measure between 8 and 18 nm in width along the same filament (Fig. 4b-f; Online Resource 4). The donor brain for the tissue used for these tomograms (Fig. 1a) had a high level of Alzheimer’s disease neuropathologic change including a severe burden of tau and amyloid-β in the absence of TDP-43 or α-synuclein aggregates in the middle frontal cortex. Based on the dimensions of the filaments, their location and orientations within axons, our knowledge that tau is present at a high burden in the region sampled, and comparison to recently published images of tau fibrils in previously frozen brain tissue [9] and isolated extracellular vesicles (Fig. S3) [8], these long twisting filaments are most likely tau aggregates. We attempted subtomogram averaging of the filaments but were unable to obtain a reliable average due to low contrast, the presence of platinum contamination on the surface of this particular lamella (this platinum surface contamination was not present on all lamella preparations), and relatively few cross-sectional views of the filaments which were noted to allow for subtomogram averaging in other samples [9].

One benefit of our approach is the ability to image myelin sheaths in a native context. Central nervous system myelin is an oligodendrocyte wrapping its processes around a neuronal axon to allow for saltatory conduction of action potentials [14, 22]. Myelination is a complex event that requires specialized signals and structures [22] thus cannot be reliably reproduced in cell culture. With our sample, even in two-dimensional cryo-EM projection images we visualize interesting myelin structures including the inner tongue of oligodendrocytes (Fig. S2c) which is thought to be important for continued myelin synthesis [22]. Future targeted imaging may allow these important structures to be comprehensively studied at higher resolution to increase our understanding of myelin structure and function.

Tomograms reveal the three-dimensional architecture of myelin, including the curvature and inter-weaving of myelin (Online Resource 3), which cannot be captured in two-dimensional projection images. Our tomograms revealed the repetitive packing of myelin around neuronal axons (Fig. 3a; Fig. 4a; Fig. 5a; Online Resource 2-3). In regions of compact myelin, we see a repeated pattern of three parallel lines: two dark outside lines representing the extracellular facing phospholipid heads (dark blue or dark red in Fig. 5b) on either side of a dark and more dense internal line (light blue or light red in Fig. 5b), the major dense line, representing the two sets of intracellular facing phospholipid head groups and myelin basic protein (MBP) that holds myelin tightly compact (Fig. 4a; Fig. 5). This repeating structure of the myelin sheath is consistent with what has been seen by traditional fixed and stained EM of brain tissue.

**Fig. 5.**
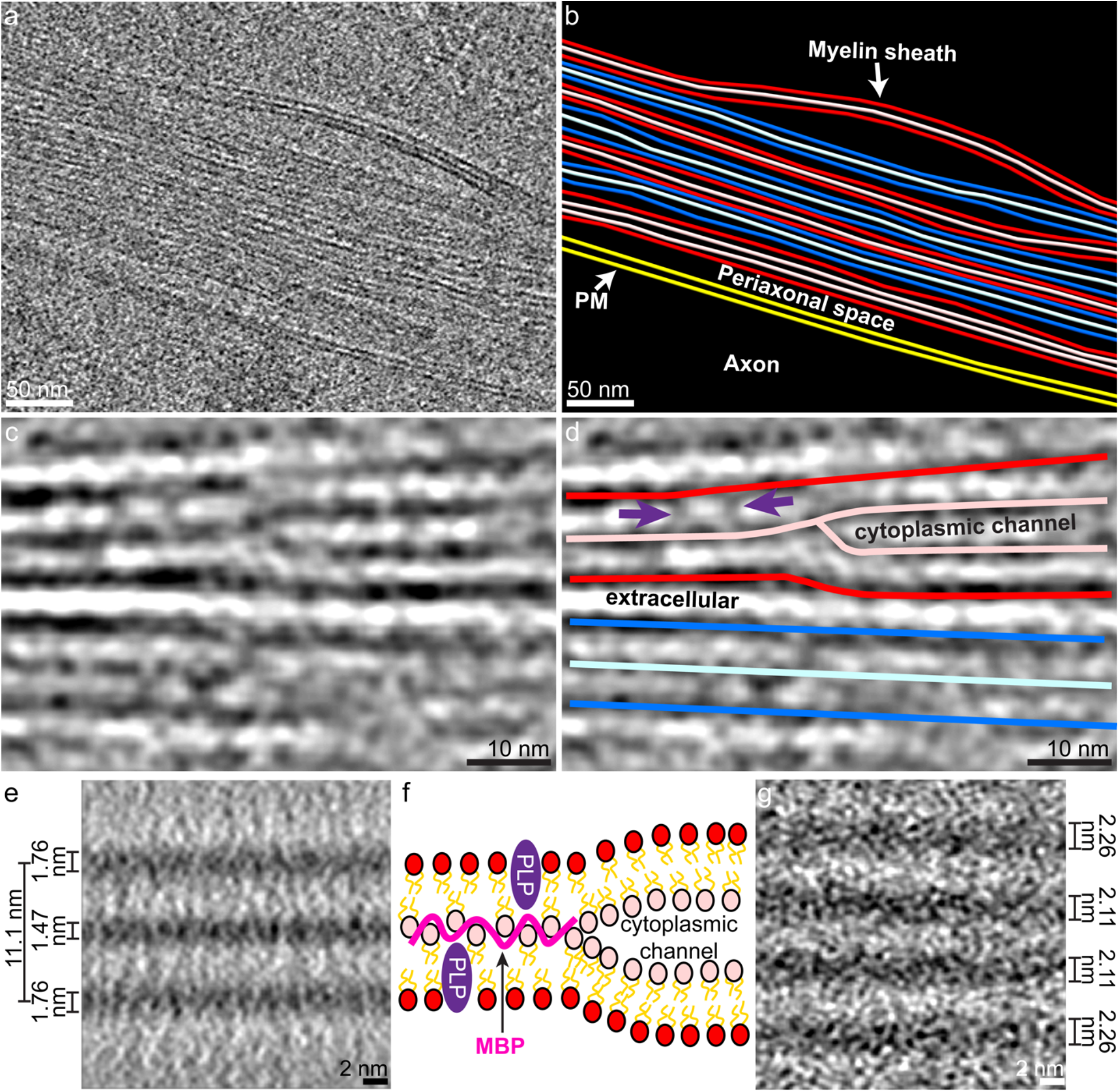
Preserved features of myelin and novel insight into myelin architecture as seen by cryo-EM in lamellae. **a** The yellow box from Fig. 4a expanded to show repeating myelination as visualized by cryo-ET. **b** segmentation of the myelination shown in **a.** Red and blue are alternated for layers of myelin with the darker shades indicating the extracellular surface of the oligodendrocyte plasma membrane and the lighter shades representing the intracellular surface of the oligodendrocyte plasma membrane. The layer of myelin closest to the axonal plasma membrane (PM; yellow) has a gap between intracellular surfaces of oligodendrocyte plasma membrane indicating an area of cytoplasmic expansion. **c** An example area of cytoplasmic expansion known as a cytoplasmic channel where the major dense line can be seen splitting into two separate electron dense lines surrounding the expanded intracellular space. **d** A tracing of the image in **c** highlighting the cytoplasmic channel and extracellular space with an adjacent compact myelin segment (the blue line is extended as a model to where density can be seen again since the myelin is changing planes). Some areas show densities within membranes (purple arrows) that may be transmembrane proteins like myelin proteolipid protein (PLP). **e** Subtomogram average of compact myelin showing three electron dense lines (average of 5 central Z slices with 2.675 Å voxel size). The outer lines are the extracellular-facing surfaces of the oligodendrocyte plasma membrane and the central line is the major dense line (the intracellular surfaces of the oligodendrocyte plasma membrane). The total width of the myelin repeating unit measures about 11.1 nm. The major dense line measures 1.47 ± 0.03 nm thick, which is less than the extracellular-facing polar head group densities that measure 1.76 ± 0.02 nm thick when averaged together. **f** Schematic of myelin architecture showing the two sets of lipid bilayers with myelin proteolipid protein (PLP) (purple) and myelin basic protein (MBP) (pink). MBP is shown tightly binding to the intracellular facing polar head groups to rigidify and compact the two sets of intracellular polar head groups in areas of compact myelin. MBP’s absence is marked by the presence of a cytoplasmic channel. **g** Subtomogram average of a cytoplasmic channel showing four electron dense lines (average of 30 central Z slices with 2.675 Å voxel size). The two outer lines are extracellular polar head groups and in two inner lines are intracellular polar head groups that compress to form the major dense line seen in **e**.

In addition to the relatively consistent compact myelin, we see areas of small cytoplasmic expansion corresponding to structures known as cytoplasmic channels (Fig. 5c-d). Cytoplasmic channels are areas of structured oligodendrocyte cytoplasmic expansion where 2’,3’-cyclic nucleotide 3’-phosphodiesterase (CNP) associates with actin to counteract MBP allowing for normal intracellular transport from the oligodendrocyte cell body to the periaxonal space [33] since intracellular transport cannot efficiently cross orthogonally through compact myelin. These cytoplasmic channels have previously only been visualized by transmission EM using relatively low-resolution techniques (HPF then freeze substitution) in part because non-cryogenic methods of fixation and staining destroy these ultrafine structures [33]. In our images of these myelin cytoplasmic channels, we observed the central major dense line splitting into two distinct lines resulting in a four-layered myelin structure (Fig. 5c-d). This allowed us to conclude that the outer set of dark lines (dark blue and dark red in Fig. 5b, d) consists of the extracellular facing polar head groups and the inner set of dark lines (light blue and light red in Fig. 5b, d) consists of the intracellular facing polar head groups, with the space between the inner lines corresponding to the cytoplasmic channel (Fig. 5c-d, f).

From inspection of myelin, we noticed that the major dense line appeared to have a similar thickness to the lines of extracellular facing polar head groups (Fig. 5c) and at split points for cytoplasmic channels the lines did not appear thinner once the intracellular facing polar head groups were separated (Fig. 5c-d). To investigate these observations, we performed subtomogram averaging of both compact myelin (Fig. 5e) and a cytoplasmic channel (Fig. 5g). We found that in regions of compact myelin, the extracellular facing polar head groups measured 1.76 ± 0.02 nm wide. Interestingly, the major dense line, representing two sets of intracellular polar head groups and MBP measured 1.47 ± 0.03 nm – less than the single sets of extracellular facing polar head groups (p=4×10^-7^; Fig. S4a). Furthermore, the subtomogram average of the region of a cytoplasmic channel showed that once the intracellular facing polar head groups split, they measured similar in width to the extracellular facing polar head groups in the same region (p=0.33; Fig. S4b). These measurements from subtomogram averaging indicate that the presence of MBP pulls the intracellular polar head groups tightly against each other. It has been previously shown that MBP is necessary for creating compact myelin [33]. It has also been suggested that as a very basic protein, MBP acts by tightly binding to the intracellular polar head groups of the oligodendrocyte plasma membrane and also that the polar head groups may serve to structure the intrinsically disordered MBP [27]. We hypothesize that MBP’s tight association with lipid polar head groups works to hold the oligodendrocyte plasma membranes together, forcing out cytoplasm and reducing vertical motion of lipids within the membrane (Fig. 5f). Extracellular facing phospholipids and the intracellular phospholipids found in cytoplasmic channels are not constrained by interactions with MBP so will have more vertical motion, thereby increasing the apparent thickness of these layers of polar head groups compared to that of the MBP bound polar head groups in subtomogram averages (Fig. 5e; Fig. S4). Based on the major dense line’s thickness of about 1.47 ± 0.03 nm, MBP likely poses as an elongated or flattened structure to make a tight mortar between the two sets of intracellular polar head groups (Fig. 5f). Future structural and biophysical studies may reveal the true structure or pattern of MBP between the oligodendrocyte plasma membranes to form compact myelin.

Beyond our indirect observations of MBP, we observed densities that appear to span the myelin membrane layer (purple arrows Fig. 5d). Myelin is known to have several transmembrane proteins including myelin proteolipid protein (PLP) (Fig. 5f). These densities that span across the lipid membrane bilayer may represent PLP and other transmembrane structural proteins that have been difficult to resolve via *in vitro* cryo-EM. Higher resolution and increased sampling will be required for subtomogram averaging in order to more definitively identify and structurally characterize these proteins.

## Discussion

Here we present the first cryo-EM visualization of native human brain tissue from unfixed, never previously frozen samples acquired at time of autopsy. We also present a method for generating vitreous human brain tissue via plunge-freezing as opposed to HPF and a method for efficiently generating on-grid lamellae at variable height using PFIB milling.

Previously, EM of human tissue has required fixation at time of autopsy followed by sectioning and staining for visualization of features, a process that introduces many artifacts and limits resolution of imaging. Many human diseases and processes, particularly those of the brain such as Alzheimer’s disease and myelination, lack high fidelity *in vitro* or animal models. Cryo-FIB milling followed by cryo-ET, while higher fidelity than traditional EM techniques, has mostly been performed on isolated cells in culture and has only recently moved towards imaging of large tissue samples [2, 12]. Most large samples have been vitrified via HPF, which has lower throughput than plunge-freezing (∼4 grids/hour with HPF [12] vs. ∼20 grids/hour with plunge-freezing), frequently uses organic solvents that need to be sublimed after freezing, can embed samples in a block of ice, and requires technical expertise and equipment modification to freeze samples directly onto a cryo-EM grid. While plunge-freezing thick samples has been shown previously [2], our results indicate that different tissue samples require different protocols to allow vitreous plunge-freezing likely due to differences in lipid and water content. We anticipate our protocol may need additional adjustments to suit other tissue types.

Large tissue samples also have been challenging for cryo-FIB milling due to milling rates. Gallium as an ion source for milling is difficult to focus at high beam currents and has been shown to interact with samples to alter their original structure [17], likely due to gallium’s depth of sample interaction. Xenon plasma, however, has comparable beam characteristics to gallium at low currents, such as beam width at a focal point, but is significantly easier to focus at high currents and has a reduced interaction depth compared to gallium making it suitable for large volume samples that would take several days to thin by gallium-based FIB milling. Our protocol highlights how currents previously only used for material science applications can make large volume FIB milling possible for biological specimens.

With this method, we generated thin lamellae in which we could already visualize the heterogeneous multicellular landscape of human brain tissue and subcellular structures within this complex milieu. We were able to identify a diverse array of subcellular compartments including putative endosomes, lysosomes, autophagic vesicles, granular vesicles, and phagophore-like structures. We also saw large proteinaceous structures *in situ*, including potential tau aggregates, that have, until recently, relied on sequential fractionation to extract and image *ex vivo*. Imaging of unfixed, never previously frozen tissue may allow for high-resolution confirmation of native tau aggregate structures as well as visualization and identification of the many different proteins that interact with tau aggregates in diseased brains. Our images also show the complex interweaving of myelin layers around an axon, a phenomenon that is not easily recapitulated *in vitro*, and we visualize areas key to normal myelin function. Future imaging of native tissue can describe *in situ* complex junctions such as the inner or outer tongue, nodes of Ranvier, or even areas of demyelination. We were also able to visualize the major dense line of myelin and cytoplasmic channels at high enough resolution to determine that the two sets of intracellular polar head groups and MBP actually pack tighter than a single set of extracellular polar head groups or a single set of intracellular polar head groups in cytoplasmic channels, thereby further elucidating how MBP may work to keep myelin compact *in vivo*.

Development of approaches, such as ours, that can attain high resolution with minimal artifacts has been recognized to be of great value. Previous work has shown subtomogram averaging of disease aggregates directly in human brain tissue via cryo-ET [9]. That study obtained novel subtomogram averages of *in situ* tau fibrils, which raise intriguing questions about differences between *in situ* and *ex vivo* tau fibril averages [9, 31]. Benefits of our workflow include use of never previously frozen brain tissue that should have better preserved cellular architecture, protein localization, and protein structure. Additionally, our method avoids use of cryo-ultramicrotome sectioning to prevent cut artifacts and distortions. However, fresh, never previously frozen tissue has drawbacks including variable postmortem interval and perimortem events, unknown pathology type and localization at time of freezing, and low availability of uncommon disease variants. Both approaches are valuable for furthering our understanding of diseases and basic biologic processes in a native context.

These initial tomograms were obtained on lamellae that were ∼400nm thick as proof of principle that human tissue can be vitrified, cryo-FIB milled, and imaged via cryo-ET. However, thickness somewhat limited ultrafine resolution as it limited the number of electrons that can be detected before radiation damage occurs. As a result, images had relatively low contrast that made data collection, image alignment, and structure visualization challenging. With further refinement of milling and imaging techniques, future tissue processing will focus on generating thinner lamellae and enhancing tilt series contrast in order to enhance resolution to allow for visualization of specific cargo and proteins *in situ*.

Our method provides a framework for imaging human disease in native environments that are not easily recapitulated in cell culture or animal models. This approach should be applicable to tissue thicknesses up to about 300 μm which is limited only in our experience by the height of the cartridge ring used for automatic loading onto a Titan Krios. Our method could also be combined with existing techniques for correlative light and electron microscopy to allow targeting of rare structures within tissue samples. While we present work in human brain tissue, this framework should be applicable to other human tissue, bulk tissue, or organoid samples.

This method is an important step towards high-resolution cryo-EM imaging of both normal and diseased human tissue in the most native context possible.

## Methods

### Human tissue

Post-mortem brain tissue was obtained at time of autopsy from brains donated to the Center for Neurodegenerative Research (CNDR) Brain Bank at the University of Pennsylvania. The illustrative cryo-ET sample highlighted in this manuscript was obtained from an 84 year-old man with a 13 year history of behavioral variant frontotemporal dementia. Neuropathologic examination revealed a high level of Alzheimer’s disease neuropathologic change (A3, B3, C3), Braak VI, moderate cerebral amyloid angiopathy, and severe arteriosclerosis. Severe tau and amyloid burden were seen in the middle frontal cortex. No TDP-43 or α-synuclein aggregates were seen.

### Sample preparation

Human brain tissue was acquired at time of autopsy from the middle frontal cortex. Tissue was acquired from autopsies performed between 7 and 28 hours postmortem. After acquisition, tissue was kept on ice. Sections of cortex less than 3 mm wide were isolated using pre-cooled razor blades on ice. Tissue sections were embedded in cool 6% low-melting point agarose (ThermoFisher, Waltham, MA) in DPBS. Tissue embedded in agarose was cooled on ice for 15 min. The agarose-embedded tissue was then superglued to vibratome stage and allowed to dry on ice for at least 15 min. Next, the vibratome stage with attached tissue was inserted into the vibratome (Model G Oxford Vibratome or Leica VT1200S, Leica Biosystems, Deerfield, IL, USA), surrounded by fresh ice, and then filled with DPBS. Tissue was cut with the vibratome set to 100 μm sections using sapphire blade (Ted Pella, Redding, CA, USA), maximum amplitude, speed 2 (Oxford) or 0.5 mm/s (Leica), and blade angle slightly below 0° (Oxford) or ∼15° as measured by vibratome guide markings (Leica). As tissue sections were cut, they were moved to cold DPBS then cold 20% glycerol with 1M trehalose in DPBS solution for 70 min. Tissue sections were then placed onto 200 mesh or 100 mesh bare copper grids (Electron Microscopy Sciences, Hatfield, PA, USA), manually blotted from behind the grid, and plunge-frozen into a liquid ethane/propane mixture using a Leica GP2 (Leica Microsystems, Deerfield, IL, USA). Frozen grids were then clipped into autogrids (ThermoFisher).

### FIB milling

Xenon plasma FIB milling was performed on a Tescan S8000X FIB/SEM with Leica VCT and VCM operating at 30 keV beam energy. The stage was cooled to cryogenic conditions. Prior to loading grids onto the FIB/SEM, the beams were manually checked and tuned on silicon. Gas injection system organometallic platinum was warmed to 70 °C. Gas injection system platinum was expelled for two min prior to loading sample grids. Grids were transferred to the FIB/SEM using a Leica VCT and VCM under cryogenic conditions. The SEM beam was used to assess grid quality and orientation prior to platinum coating. Grids were oriented so the FIB beam would be close to orthogonal to horizontal grid bars. After initial assessment, grids were coated with gas injection system platinum for 1 min total in 15 s increments with 15 s rest intervals between injections. This initial gas injection system platinum deposition was carried out at −30° rotation, 0° tilt. The organometallic platinum layer was then cured by rastering the FIB beam over the full tissue sample three times using 10 nA, 30 keV at field of view (FoV) 1200.7 μm with raster speed 4 at −30° rotation, 10° tilt.

Trench milling was performed to reveal an internal tissue face. Trench milling was performed at −30° rotation, 5° tilt, with the FIB nearly orthogonal to the plane of the grid (15° being orthogonal). Initial milling steps at 1 μA and 100 nA used box milling patterns, and all milling steps below 100 nA used directional polishing patterns. Milling was initiated at 1 μA current until about 80-100 μm from the targeted finishing location of trench milling. Targeted trench finishing was located just behind a grid bar to allow for maximum imaging area and prevent grid bars from occluding transmission EM imaging. 100 nA polishing pattern was used to mill up to the grid bar directly in front of future lamellae positions. 100 nA square pattern was then used to mill away grid bars in front of future lamellae positions. 10 nA current with polishing pattern was used to smooth curtaining at all spots of potential lamellae.

After trench milling, gas injection system organometallic platinum was deposited onto the exposed tissue face with a retracted gas injection system needle at −165° rotation, 40° tilt (gas injection system needle orthogonal to the tissue face). Curing of this platinum layer using the FIB beam was performed at −150° rotation, 20-27° tilt depending on field of view so that all areas of platinum covering tissue to be milled were cured. Curing was performed as described above. Gas injection system platinum was deposited for a 5 minute in total, with iterative depositions alternating with curing by the following pattern: 15 s deposition, cure, 15 s deposition, cure, 30 s deposition, cure, 30 s deposition, cure. 30 s deposition, cure, 1 min deposition, cure, 1 min deposition, cure, 1 min deposition, cure.

Lamellae were milled at −150° rotation with a final tilt angle of between 19° and 23°. Expansion joints were made on either side of the lamellae using 10 nA beam to roughly 8 μm wide. The 10nA beam was also used to thin the tissue to a thickness of 10-20um. Material was first removed from below the intended lamella position from an initial tilt angle of 21-27°. The tilt angle was then reduced incrementally when possible as material was removed from the underside of the tissue. After excess material from beneath the final lamella was removed, the milling tilt angle was adjusted to −1° relative to the final intended lamella angle (i.e. if the final lamella was intended to be 19°, then milling from below was performed at 18°). Material was removed above the intended lamella position at +1° from the final intended lamella angle, also using the 10 nA beam. Starting at 10-20 μm thickness, over and under tilting was reduced to +/- 0.5° and the beam current was reduced to 1 nA. Around 1-3 μm thickness, over and under tilting was reduced to +/- 0.3° and the beam current was reduced to 100 pA. Final polishing steps were carried out at 100 pA and then 50 pA. This final polishing step was carried out on all lamellae on a given grid just prior to the removal of the grid from the FIB/SEM, to ensure minimal surface ice deposition. The lamellae were polished to a final thickness of <400 nm as measured by FIB images. Lamella grids were unloaded under cryogenic conditions using Leica VCT and VCM and stored in liquid nitrogen until they were imaged via transmission EM.

### Cryo-EM

Cryo-EM was performed on a Thermo Fisher Krios G3i 300 keV field emission cryo-transmission electron microscope. Imaging was performed using the SerialEM software [21] on a K3 direct electron detector (Gatan, Pleasanton, CA, USA) operated in electron-counted mode. Imaging was performed with a slit width of 50 eV between magnifications of x4,800 and x19,500. Imaging at magnification of x33,000 was performed with a slit width of 20 eV. Imaging at magnification of x82 and x470 was used to assess grid orientation to ensure lamellae were loaded orthogonal to tilt axis and assess suitability for imaging after transfer. Further montages were taken at magnification x4,800 and x19,500 to identify features in lamellae that would be suitable for high magnification imaging. Tilt series were collected with a span of 72° (+/- 36° from the tilt angle at which lamellae were orthogonal to electron beam; dose-symmetric scheme) with 3° increments at magnification of x33,000 (with a corresponding pixel size of 2.67 Å) and a defocus target of −6 μm. Tilt series were collected with a cumulative dose of between 86 and 97 e^-^ Å^-2^.

### Image Processing

Montages at magnification of x19,500 were rotated to align curtaining with the vertical axis of the image. A vertical filter was applied with 95% tolerance of direction in Fiji [29] to remove curtaining artifacts.

Tilt series were first filtered in Fiji using a high-pass filter of 1000 pixels and vertical filter with 95% tolerance of direction. Tilt series were aligned using platinum deposition on the surface of the lamellae as fiducials and reconstructed into tomograms via weighted back projection in IMOD [16]. Tomograms were reconstructed with an exact filter of 90 unbinned pixels or a SIRT-like filter of 10 iterations. For visualization, some tomograms were denoised using 3D denoising with or without a gaussian filter in Topaz [3].

For subtomogram averaging of compact myelin, model points were placed along straight regions of the major dense line using IMOD in 4x binned (10.70 Å voxel size) tomograms. Additional model points were added every 2 voxels using ‘addModPts’ command in PEET [24] for a total of 4043 model points. PEET was used for subtomogram averaging in unbinned tomograms (2.675 Å voxel size) with iterations of rotational and translational searching to align the different regions of myelin with a box size of 96x96x96. Fiji was used to define values of the average intensity for five adjacent central sets of 10 sequential voxels in Z of the subtomogram average ranging from Z=20 to Z=70. From these values, full width half maximum was used to measure the width of the major dense line and extracellular facing polar head groups by fitting the voxel intensity over each line to a gaussian curve. In the average of compact myelin, the two outer lines were grouped together. A Student’s t-Test, with a two-tailed distribution was used to determine significance of the full width half maximum measurement for inside dark lines compared to outside dark lines. A similar method was employed for subtomogram averaging of cytoplasmic channels and subsequent measurement of polar head group density with model points being placed at the center of a cytoplasmic channel for a total of 127 model points. For the cytoplasmic channel average, the two outer lines were grouped together and the two inner lines were grouped together. Width of the total myelin repeating unit was measured from maximum to maximum of the extracellular facing lipid head densities.

Manual segmentation/tracing of myelin components were produced in IMOD and imported into ChimeraX [26] for visualization and figure generation.

## Supporting information

Online Resource 1

Online Resource 2

Online Resource 3

Online Resource 4

## Acknowledgements

We are grateful to the patients, families and caregivers that make this research possible. We thank Jamie Ford at the Singh Center for Nanotechnology and Stefan Steimle at the Beckman Center for Cryo-Electron Microscopy at the University of Pennsylvania for their assistance. The authors gratefully acknowledge use of facilities and instrumentation supported by the National Science Foundation through the University of Pennsylvania Materials Research Science and Engineering Center (DMR-2309043). This study was supported by the DeCrane Family Fund for PPA Research and grants RF1AG065341 to E.B.L. and Y.-W.C., P30AG072979, P01AG066597, and U19AG062418 from the National Institute of Health to E.B.L.; a David and Lucile Packard Fellowship for Science and Engineering (2019-69645), Burroughs Wellcome Fund Investigators in the Pathogenesis of Infectious Disease Program (1022785), and a Pennsylvania Department of Health FY19 Health Research Formula Fund to Y.-W.C.; and R35GM130302 from the National Institute of Health to B.E.B. Training support was provided by the National Institute of Health T32GM132039 and F30AG077756 to B.C.C., and F30CA261198 to K.K.

## Author contributions

BCC, KK, EBL, and Y-WC conceived the study and designed the experiments. BCC and KK performed the experiments. BCC, KK, EBL, and Y-WC analyzed the data. BEB provided critical input on the study. The manuscript was written by BCC, KK, EBL, and Y-WC with input from all the authors.

## References

1. Arseni D, Hasegawa M, Murzin AG, Kametani F, Arai M, Yoshida M, Ryskeldi-Falcon B (2022) Structure of pathological TDP-43 filaments from ALS with FTLD. Nature 601:139–143. doi: 10.1038/s41586-021-04199-3

2. Bäuerlein FJB, Renner M, Chami DE, Lehnart SE, Pastor-Pareja JC, Fernández-Busnadiego R (2023) Cryo-electron tomography of large biological specimens vitrified by plunge freezing. bioRxiv. doi: 10.1101/2021.04.14.437159

3. Bepler T, Kelley K, Noble AJ, Berger B (2020) Topaz-Denoise: general deep denoising models for cryoEM and cryoET. Nat Commun 11:5208. doi: 10.1038/s41467-020-18952-1

4. Berger C, Dumoux M, Glen T, Yee NB -y, Mitchels JM, Patáková Z, Darrow MC, Naismith JH, Grange M (2023) Plasma FIB milling for the determination of structures in situ. Nat Commun 14:629. doi: 10.1038/s41467-023-36372-9

5. Bieber A, Capitanio C, Erdmann PS, Fiedler F, Beck F, Lee C-W, Li D, Hummer G, Schulman BA, Baumeister W, Wilfling F (2022) In situ structural analysis reveals membrane shape transitions during autophagosome formation. Proc Natl Acad Sci 119:e2209823119. doi: 10.1073/pnas.2209823119

6. Burneg TL, Kelley R, Winiarski B, Contreras L, Daly M, Gholinia A, Burke MG, Withers PJ (2016) Large volume serial section tomography by Xe Plasma FIB dual beam microscopy. Ultramicroscopy 161:119–129. doi: 10.1016/j.ultramic.2015.11.001

7. Cheville NF, Stasko J (2014) Techniques in Electron Microscopy of Animal Tissue. Vet Pathol 51:28–41. doi: 10.1177/0300985813505114

8. Fowler SL, Behr TS, Turkes E, Cauhy PM, Foiani MS, Schaler A, Crowley G, Bez S, Ficulle E, Tsefou E, O’Brien DP, Fischer R, Geary B, Gaur P, Miller C, D’Acunzo P, Levy E, Duff KE, Ryskeldi-Falcon B (2023) Tau filaments are tethered within brain extracellular vesicles in Alzheimer’s disease. bioRxiv. doi: 10.1101/2023.04.30.537820

9. Gilbert MAG, Fatima N, Jenkins J, O’Sullivan TJ, Schertel A, Halfon Y, Morrema THJ, Geibel M, Radford SE, Hoozemans JJM, Frank RAW (2023) In situ cryo-electron tomography of β-amyloid and tau in post-mortem Alzheimer’s disease brain. bioRxiv. doi: 10.1101/2023.07.17.549278

10. Han H-M, Zuber B, Dubochet J (2008) Compression and crevasses in vitreous sections under different cupng conditions. J Microsc 230:167–171. doi: 10.1111/j.1365-2818.2008.01972.x

11. Hildebrand C, Remahl S, Persson H, Bjartmar C (1993) Myelinated nerve fibres in the CNS. Prog Neurobiol 40:319–384. doi: 10.1016/0301-0082(93)90015-K

12. Kelley K, Raczkowski AM, Klykov O, Jaroenlak P, Bobe D, Kopylov M, Eng ET, Bhabha G, Poger CS, Carragher B, Noble AJ (2022) Waffle Method: A general and flexible approach for improving throughput in FIB-milling. Nat Commun 13:1857. doi: 10.1038/s41467-022-29501-3

13. Kelley RD, Song K, Leer BV, Wall D, Kwakman L (2013) Xe+ FIB Milling and Measurement of Amorphous Silicon Damage. Microsc Microanal 19:862–863. doi: 10.1017/S1431927613006302

14. Kinney HC, Volpe JJ (2018) Chapter 8 - Myelination Events. In: Volpe JJ, Inder TE, Darras BT, de Vries LS, du Plessis AJ, Neil JJ, Perlman JM (eds) Volpe’s Neurology of the Newborn (Sixth Edition). Elsevier, pp 176–188

15. Korogod N, Petersen CC, Knog GW (2015) Ultrastructural analysis of adult mouse neocortex comparing aldehyde perfusion with cryo fixation. eLife 4:e05793. doi: 10.7554/eLife.05793

16. Kremer JR, Mastronarde DN, McIntosh JR (1996) Computer visualization of three-dimensional image data using IMOD. J Struct Biol 116:71–76. doi: 10.1006/jsbi.1996.0013

17. Lucas BA, Grigorieff N (2023) Quantification of gallium cryo-FIB milling damage in biological lamellae. Proc Natl Acad Sci 120:e2301852120. doi: 10.1073/pnas.2301852120

18. Mahamid J, Pfeffer S, Schaffer M, Villa E, Danev R, Kuhn Cuellar L, Förster F, Hyman AA, Plitzko JM, Baumeister W (2016) Visualizing the molecular sociology at the HeLa cell nuclear periphery. Science 351:969–972. doi: 10.1126/science.aad8857

19. Mahamid J, Schampers R, Persoon H, Hyman AA, Baumeister W, Plitzko JM (2015) A focused ion beam milling and liu-out approach for site-specific preparation of frozen-hydrated lamellas from multicellular organisms. J Struct Biol 192:262–269. doi: 10.1016/j.jsb.2015.07.012

20. Marko M, Hsieh C, Schalek R, Frank J, Mannella C (2007) Focused-ion-beam thinning of frozen-hydrated biological specimens for cryo-electron microscopy. Nat Methods 4:215–217. doi: 10.1038/nmeth1014

21. Mastronarde DN (2005) Automated electron microscope tomography using robust prediction of specimen movements. J Struct Biol 152:36–51. doi: 10.1016/j.jsb.2005.07.007

22. Meschkat M, Steyer AM, Weil M-T, Kusch K, Jahn O, Piepkorn L, Agüi-Gonzalez P, Phan NTN, Ruhwedel T, Sadowski B, Rizzoli SO, Werner HB, Ehrenreich H, Nave K-A, Möbius W (2022) White mager integrity in mice requires continuous myelin synthesis at the inner tongue. Nat Commun 13:1163. doi: 10.1038/s41467-022-28720-y

23. Möbius W, Nave K-A, Werner HB (2016) Electron microscopy of myelin: Structure preservation by high-pressure freezing. Brain Res 1641:92–100. doi: 10.1016/j.brainres.2016.02.027

24. Nicastro D, Schwartz C, Pierson J, Gaudege R, Porter ME, McIntosh JR (2006) The Molecular Architecture of Axonemes Revealed by Cryoelectron Tomography. Science 313:944–948. doi: 10.1126/science.1128618

25. Peters A, Palay SL, Webster H deF (1976) The Fine Structure of the Nervous System: The Neurons and Supporting Cells. W. B. Saunders Company

26. Pegersen EF, Goddard TD, Huang CC, Meng EC, Couch GS, Croll TI, Morris JH, Ferrin TE (2021) UCSF ChimeraX: Structure visualization for researchers, educators, and developers. Protein Sci Publ Protein Soc 30:70–82. doi: 10.1002/pro.3943

27. Raasakka A, Ruskamo S, Kowal J, Barker R, Baumann A, Martel A, Tuusa J, Myllykoski M, Bürck J, Ulrich AS, Stahlberg H, Kursula P (2017) Membrane Association Landscape of Myelin Basic Protein Portrays Formation of the Myelin Major Dense Line. Sci Rep 7:4974. doi: 10.1038/s41598-017-05364-3

28. Schaffer M, Pfeffer S, Mahamid J, Kleindiek S, Laugks T, Albert S, Engel BD, Rummel A, Smith AJ, Baumeister W, Plitzko JM (2019) A cryo-FIB liu-out technique enables molecular-resolution cryo-ET within native Caenorhabditis elegans tissue. Nat Methods 16:757–762. doi: 10.1038/s41592-019-0497-5

29. Schindelin J, Arganda-Carreras I, Frise E, Kaynig V, Longair M, Pietzsch T, Preibisch S, Rueden C, Saalfeld S, Schmid B, Tinevez J-Y, White DJ, Hartenstein V, Eliceiri K, Tomancak P, Cardona A (2012) Fiji: an open-source plaworm for biological-image analysis. Nat Methods 9:676–682. doi: 10.1038/nmeth.2019

30. Sele M, Wernitznig S, Lipovšek S, Radulović S, Haybaeck J, Birkl-Toeglhofer AM, Wodlej C, Kleinegger F, Sygulla S, Leoni M, Ropele S, Leitinger G (2019) Optimization of ultrastructural preservation of human brain for transmission electron microscopy auer long post-mortem intervals. Acta Neuropathol Commun 7:144. doi: 10.1186/s40478-019-0794-3

31. Shi Y, Zhang W, Yang Y, Murzin AG, Falcon B, Kotecha A, van Beers M, Tarutani A, Kametani F, Garringer HJ, Vidal R, Hallinan GI, Lashley T, Saito Y, Murayama S, Yoshida M, Tanaka H, Kakita A, Ikeuchi T, Robinson AC, Mann DMA, Kovacs GG, Revesz T, Ghep B, Hasegawa M, Goedert M, Scheres SHW (2021) Structure-based classification of tauopathies. Nature 598:359–363. doi: 10.1038/s41586-021-03911-7

32. Smith NS, Skoczylas WP, Kellogg SM, Kinion DE, Tesch PP, Sutherland O, Aanesland A, Boswell RW (2006) High brightness inductively coupled plasma source for high current focused ion beam applications. J Vac Sci Technol B Microelectron Nanometer Struct Process Meas Phenom 24:2902– 2906. doi: 10.1116/1.2366617

33. Snaidero N, Velte C, Myllykoski M, Raasakka A, Ignatev A, Werner HB, Erwig MS, Möbius W, Kursula P, Nave K-A, Simons M (2017) Antagonistic Functions of MBP and CNP Establish Cytosolic Channels in CNS Myelin. Cell Rep 18:314–323. doi: 10.1016/j.celrep.2016.12.053

34. Studer D, Humbel BM, Chiquet M (2008) Electron microscopy of high pressure frozen samples: bridging the gap between cellular ultrastructure and atomic resolution. Histochem Cell Biol 130:877–889. doi: 10.1007/s00418-008-0500-1

35. Yang Y, Arseni D, Zhang W, Huang M, Lövestam S, Schweighauser M, Kotecha A, Murzin AG, Peak-Chew SY, Macdonald J, Lavenir I, Garringer HJ, Gelpi E, Newell KL, Kovacs GG, Vidal R, Ghep B, Ryskeldi-Falcon B, Scheres SHW, Goedert M (2022) Cryo-EM structures of amyloid-β 42 filaments from human brains. Science 375:167–172. doi: 10.1126/science.abm7285

36. Yang Y, Shi Y, Schweighauser M, Zhang X, Kotecha A, Murzin AG, Garringer HJ, Cullinane PW, Saito Y, Foroud T, Warner TT, Hasegawa K, Vidal R, Murayama S, Revesz T, Ghep B, Hasegawa M, Lashley T, Scheres SHW, Goedert M (2022) Structures of α-synuclein filaments from human brains with Lewy pathology. Nature 10.1038/s41586-022-05319–3. doi: 10.1038/s41586-022-05319-3

37. Zhang T-Y, Tan P-C, Xie Y, Zhang X-J, Zhang P-Q, Gao Y-M, Zhou S-B, Li Q-F (2020) The combination of trehalose and glycerol: an effective and non-toxic recipe for cryopreservation of human adipose-derived stem cells. Stem Cell Res Ther 11:460. doi: 10.1186/s13287-020-01969-0

38. Zhao DY, Bäuerlein FJB, Saha I, Hartl FU, Baumeister W, Wilfling F (2023) Autophagy preferentially degrades non-fibrillar polyQ aggregates. bioRxiv. doi: 10.1101/2023.08.08.552291

39. Zhong X, Wade CA, Withers PJ, Zhou X, Cai C, Haigh SJ, Burke MG (2021) Comparing Xe+pFIB and Ga+FIB for TEM sample preparation of Al alloys: Minimising FIB-induced artefacts. J Microsc 282:101–112. doi: 10.1111/jmi.12983

