## Supplemental Figures for "Native ultrastructure of fresh human brain vitrified directly from autopsy revealed by cryo-electron tomography with cryo-plasma focused ion beam milling"

**Affiliations**

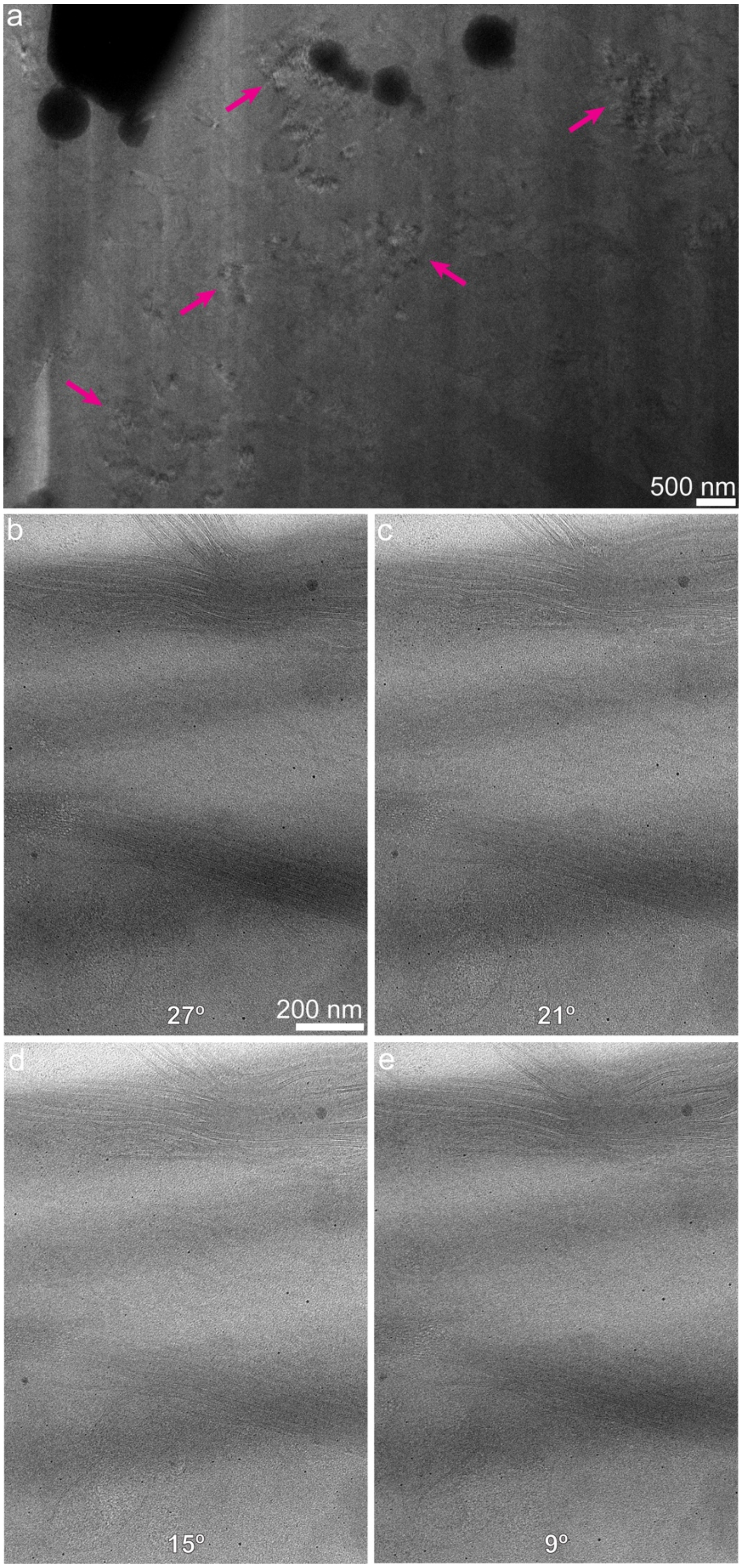


**Fig. S1** Lamellae prepared from tissue incubated with 10% glycerol show signs of crystalline ice whereas lamellae prepared from tissue incubated with 20% glycerol and 1M trehalose show no signs of crystalline ice. **a** Two-dimensional projection image of lamella from tissue frozen after 15 min incubation in 10% glycerol showing Bragg reflections indicative of crystalline ice (pink arrows). **b-e** Multiple tilt angles from a representative tilt series taken on a lamella prepared from tissue incubated with 20% glycerol and 1M trehalose showing no indication of crystalline ice within the lamella.

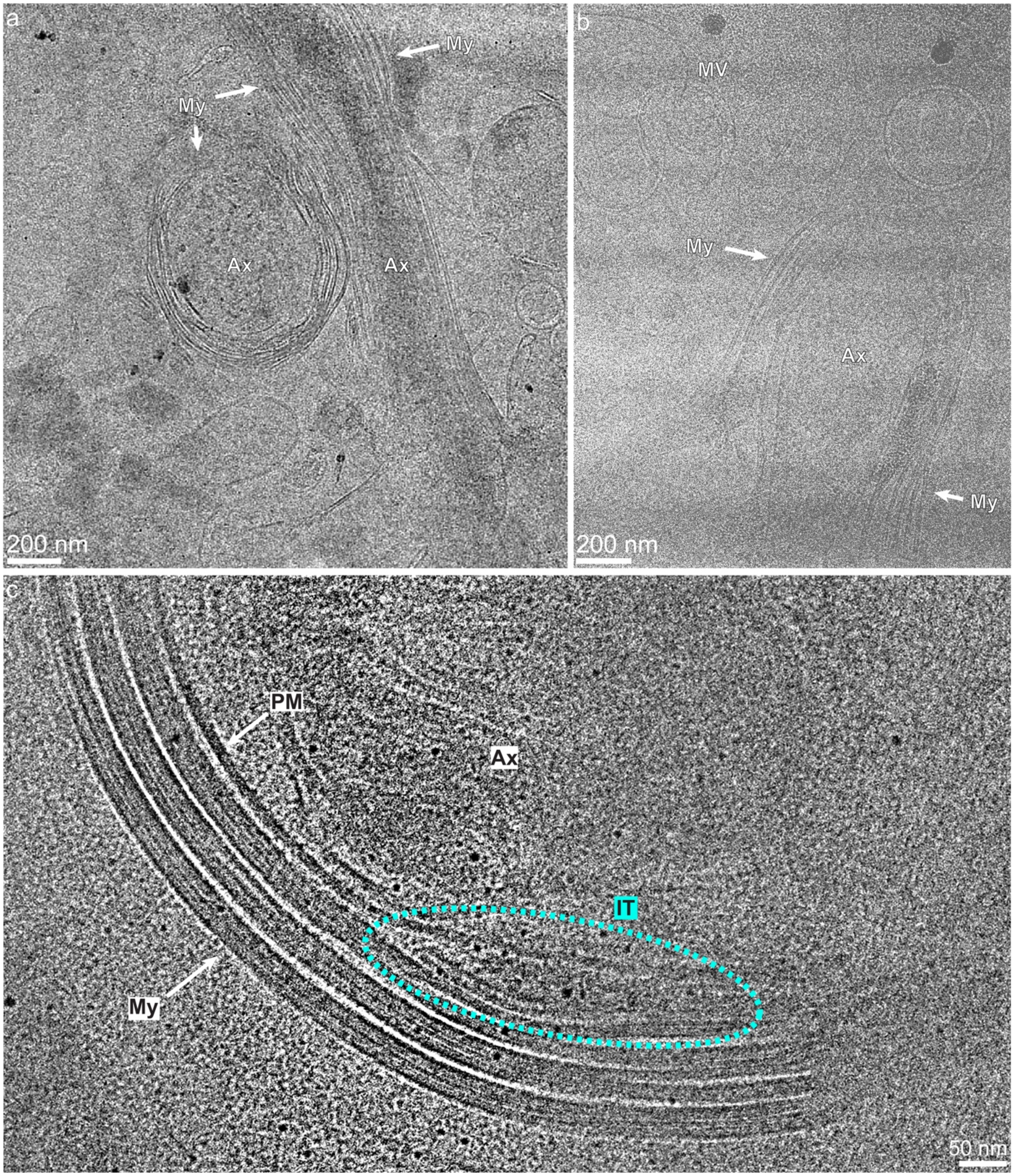


**Fig. S2** Two-dimensional projection images showing diverse cellular features within lamellae. **a-c** show images taken at x19,500 showing diverse cellular features including myelinated (My) axons (Ax), multi-layered vesicles (MV), and the inner tongue (IT) of an oligodendrocyte – a cytoplasmic expansion near the plasma membrane (PM) of an axon that is thought to be important for continued myelin generation.

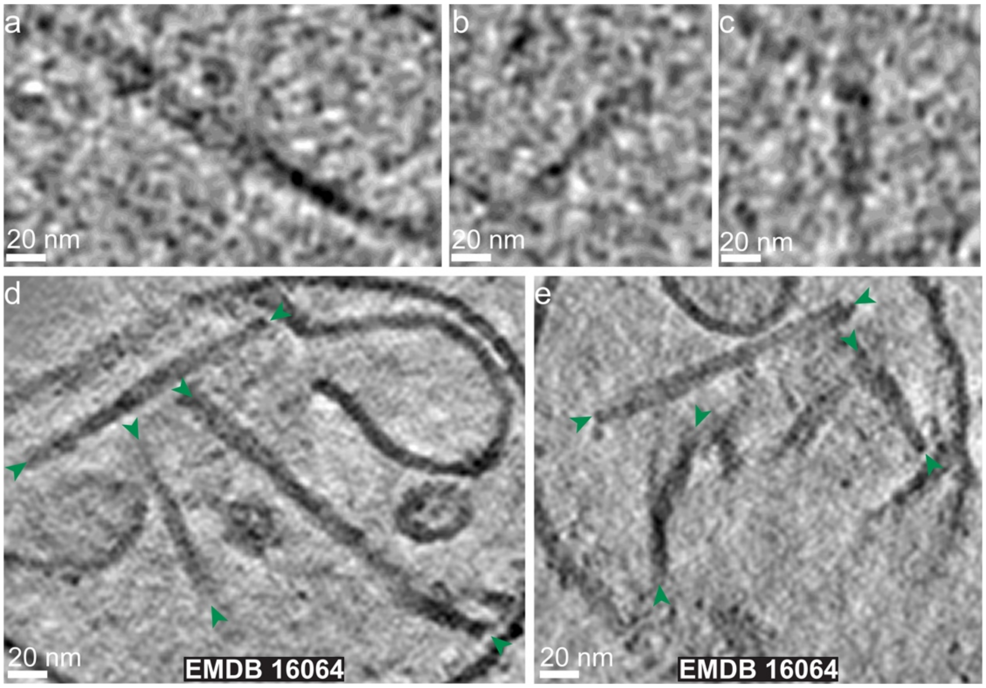


**Fig. S3** Tau fibrils seen in lamellae generated directly in human brain tissue look similar to tau fibrils seen in extracellular vesicles extracted from human brain. **a-c** Example tau fibrils from tomogram cross-section (The same images used in Fig. 4d-f). **d** and **e** are example cross-sections from tomogram of tau (green arrowheads at ends of fibrils) within extracellular vesicles purified from human brain (EMDB:16064) [8]

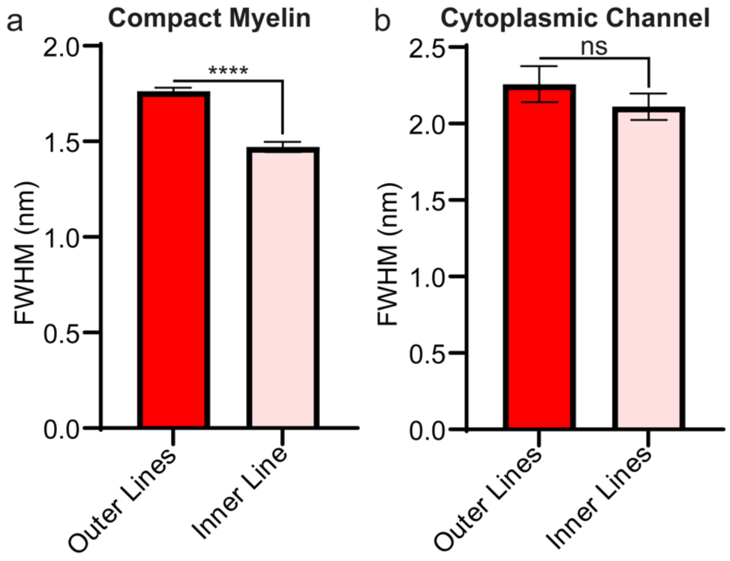


**Fig. S4** Full width half maximum (FWHM) measurements of subtomogram averages of compact myelin and cytoplasmic channel myelin. **a-b** FWHM from average pixel intensity of five adjacent regions of ten Z slices in subtomogram averages of compact myelin (**a**) and cytoplasmic channel myelin (**b**) (points are mean ± SEM; Student's t-Test: ****$p<0.0001$).

**Online Resource 1** A movie panning through slices of the tomogram shown in Figure 2d-e with an autophagic vesicle and granular vesicles labeled. Scale bar is 100 nm.

**Online Resource 2** A movie panning through the slices of the tomogram shown in Figure 3 with examples of granular vesicles, autophagic vesicles, axons (myelinated and unmyelinated), and phagophore-like structures (Ph) labeled. Scale bar is 100 nm.

**Online Resource 3** A movie panning through the slices of the tomogram shown in Figure 4 and Figure 5a, c showing a myelinated axon with potential tau fibrils running parallel to the axon. Scale bar is 100 nm.

**Online Resource 4** A movie of the same tomogram from Online Resource 3, Figure 4, and Figure 5a, c with three-dimensional denoising and gaussian filtering to better visualize potential tau fibrils running parallel to the myelinated axon. Examples of potential tau fibrils within the tomogram are labeled. Scale bar is 100 nm.
